# Active licking shapes cortical taste coding

**DOI:** 10.1101/2022.05.13.491862

**Authors:** Camden Neese, Cecilia G. Bouaichi, Tom Needham, Martin Bauer, Richard Bertram, Roberto Vincis

**Affiliations:** Florida State University, Department of Statistics; Florida State University, Department of Mathematics; Florida State University, Department of Mathematics and Programs in Neuroscience and Molecular Biophysics; Florida State University, Department of Biological Science and Program in Neuroscience

## Abstract

Neurons in the gustatory cortex (GC) represent taste through time-varying changes in their spiking activity. The predominant view is that the neural firing rate represents the sole unit of taste information. It is currently not known whether the phase of spikes relative to lick timing is used by GC neurons for taste encoding. To address this question, we recorded spiking activity from >500 single GC neurons in male and female mice permitted to freely lick to receive four liquid gustatory stimuli and water. We developed a set of data analysis tools to determine the ability of GC neurons to discriminate gustatory information and then to quantify the degree to which this information exists in the spike rate versus the spike timing or phase relative to licks. These tools include machine learning algorithms for classification of spike trains and methods from geometric shape and functional data analysis. Our results show that while GC neurons primarily encode taste information using a rate code, the timing of spikes is also an important factor in taste discrimination. A further finding is that taste discrimination using spike timing is improved when the timing of licks is considered in the analysis. That is, the interlick phase of spiking provides more information than the absolute spike timing itself. Overall, our analysis demonstrates that the ability of GC neurons to distinguish among tastes is best when spike rate and timing is interpreted relative to the timing of licks.

**Significance Statement:** Neurons represent information from the outside world via changes in their number of action potentials (spikes) over time. This study examines how neurons in the mouse gustatory cortex (GC) encode taste information when gustatory stimuli are experienced through the active process of licking. We use electrophysiological recordings and data analysis tools to evaluate the ability of GC neurons to distinguish tastants and then to quantify the degree to which this information exists in the spike rate versus the spike timing relative to licks. We show that the neuron’s ability to distinguish between tastes is higher when spike rate and timing are interpreted relative to the timing of licks, indicating that the lick cycle is a key factor for taste processing.

## Introduction

Our motivation to eat depends on the taste of food and on the reward experienced while eating. Taste information is processed through neural computations that occur in interconnected brain areas that include the gustatory portion of the insular cortex (GC), the primary cortical area responsible for processing taste information (Vincis and Fontanini, 2019; Spector and Travers, 2005). Neurons of the GC have been extensively studied and are known to represent taste through changes in their spiking activity. These responses reflect the ongoing processing and integration of diverse gustatory information. For instance, GC neurons respond to the chemosensory qualities and hedonic value of gustatory stimuli while also accounting for sensory cues that anticipate the availability of food, and behavioral states (Katz et al., 2001; Samuelsen et al., 2012; Vincis and Fontanini, 2016; Fontanini and Katz, 2006). The predominant view is that the changes in neural firing rate over seconds-long temporal windows represents the basic unit of gustatory information through which features of taste stimuli, such as their identity and hedonic value, are extracted (Katz et al., 2001; Levitan et al., 2019; Jezzini et al., 2013; Bouaichi and Vincis, 2020).

Key questions regarding the dynamics of taste-evoked spiking activity in single cortical neurons remain unanswered. Liquid gustatory stimuli are sensed by rodents through stereotyped licking behavior (Travers et al., 1997). This process, much like the role of sniffing for the sense of smell (Shusterman et al., 2011), provides rhythmic oro-facial motor activity to the brain that can be used as a metronome against which neural activity can be aligned. Yet, it is currently unknown whether the lick cycle plays a role in taste coding. How important is the rate of spikes within “lick-cycle-long” time intervals (as opposed to rate changes over randomly defined and lick-unrelated temporal windows) in GC neurons for distinguishing tastes? The second question involves action potential timing between licks. Does the time of spiking relative to licks (phase coding) contain information that can be used by GC neurons to discriminate different stimuli? Finally, if phase coding is used by GC coding neurons, how much does it improve taste discrimination?

To address these questions, we recorded spiking activity from single GC neurons in mice permitted to freely lick to receive four liquid gustatory stimuli (sucrose, NaCl, citric acid and quinine) and water (Bouaichi and Vincis, 2020). We then used a supervised machine learning algorithm, called a support vector machine, and methods from shape and functional data analysis to determine the extent to which taste information can be distinguished using spike rate, spike timing, or both. We then performed a separate analysis in which spike timing was measured with respect to the timing of licks, converting spike timing to spike phase. We observed that while the spike rate is the dominant factor for coding taste information, temporal information also contributes. Finally, the latter contribution is greater when the spike timing is relative to the timing of licks. Thus, the ability to distinguish between tastes is improved when spike rate and timing are interpreted relative to the timing of licks.

## Materials and Methods

### Data acquisition

The experiments in this study are performed on 12 wild type C57BL/6J adult mice (10-20 weeks old; 6 males and 6 females). Mice are purchased from The Jackson Laboratory (Bar Harbor, ME); upon arrival, mice are housed on a 12h/12h light-dark cycle and had ad-libitum access to food and water. Experiments and training are performed during the light portion of the cycle.

The experimental data set consists of 529 neurons; 283 neurons come from a previously published data set (Bouaichi and Vincis, 2020) while 246 neurons come from additional recordings. It is important to note that there are no differences in experimental conditions between the two data sets. Six days before training began, mice are water restricted and maintained at 85% of their pre-surgical weight. All experiments are reviewed and approved by the Florida State University Institutional Animal Care and Use Committee (IACUC) under protocol “PROTO202100006”. Prior to surgery, mice are anesthetized with a mixture of ketamine/dexmedetomidine (70 mg/kg; 1 mg/kg). The depth of anesthesia is monitored regularly via visual inspection of breathing rate, whisker reflexes, and by periodically assessing (every 30 minutes) the tail reflex. Anesthesia is supplemented by 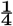 the original dose of ketamine as needed throughout the surgery. A heating pad is used to maintain body temperature at 35°C. At the start of surgery, mice are also dosed with dexamethasone (1-4 mg/kg) and lidocaine or bupivacaine HCl 2% with epinephrine (2-4 mg.kg diluted to 0.5% solution, s.c.). In addition, lactate solutions are administered every 0.5 h during surgery at volumes of 0.5 ml. After the achievement of surgical level of anesthesia, the animal’s head is cleaned, disinfected (with iodine solution and 70% alcohol), and finally fixed on a stereotaxic holder. A small craniotomy is made over the gustatory cortex (GC, relative to bregma: +1.2 AP;+3.6 or −3.6 ML) and a movable array of 8 tetrodes and one single reference wire (Sandvik-Kanthal, PX000004 with a final impedance of 200-300 kohm for tetrodes and 20-30 kohm for the reference wire) are slowly (over 10-15 minutes) lowered to reach a final position 100-150 μm dorsal to GC and fixed on the skull using dental acrylic; they are further lowered 200 μm before the first day of recording and 80 μm after each recording. Next, a small and light head-bolt is cemented to the acrylic head cap for restraint. In addition, a second craniotomy is then drilled on top of the visual cortex where a ground wire (A-M system, Cat. No. 781000) is lowered 300 μm below the brain surface. Before implantation, tetrode wires are coated with a lipophilic fluorescent dye (DiI; Sigma-Aldrich), allowing us to visualize the exact location of the tetrode bundle at the end of each experiment.

One week after the start of the water restriction regimen, mice are progressively habituated to head-restraint procedures. During restraining, the body of the mouse is covered with a semicircular opaque plastic shelter to constrain the animal’s body movements without stressful constriction (Figure 1A). The taste delivery system, licking detection and behavioral paradigm are described in detail in (Bouaichi and Vincis, 2020). Briefly, the behavioral training and recording sessions take place within a Faraday cage (Type II 36X36X40H CleanBench, TMC) to facilitate electrophysiological recording. Mice are habituated to be head restrained for short (5 min) daily sessions and gradually progressed (over days) toward longer sessions. Following the habituation to restraint, they are trained with a fixed ratio schedule, in which the mice learn to lick a dry spout six times to trigger the delivery of 3 μl of one of four prototypical tastants: 200 mM sucrose, 50 mM NaCl, 10 mM citric acid, 0.5 mM quinine, or pure water (Figure 1C). The tastes and concentrations were chosen for multiple reasons: 1) the concentrations of all four gustatory stimuli are well above their detection threshold in mice (Boughter et al., 2005; Delay et al., 2006; Ishiwatari and Bachmanov, 2012); 2) the sour and bitter stimuli are not too aversive, so that the animals remain engaged in the task and actively lick for a substantial number of trials (at least 15 for each taste, allowing proper statistical analysis of neural data); 3) they provide compatibility with prior awake-behaving taste electrophysiology studies in mice (Levitan et al., 2019; Dikecligil et al., 2020; Graham et al., 2014); 4) they represent a broad range of taste qualities and hedonic values. The choice of using a single 3 μl droplet of fluid as stimulus originated from experimental evidence indicating that mice are capable of discriminating taste in a single 2 μl droplet (smaller than the amount used in our study) of taste solutions delivered from a licking spout (Graham et al., 2014). In addition, GC taste tuning profiles recorded in active licking mice using a single 2 (Dikecligil et al., 2020), 3 (Bouaichi and Vincis, 2020) or 6 μl (Chen et al., 2021) droplet of fluid are comparable with those obtained in a recent mouse study in which larger boli (12 μl) of taste were delivered through intraoral cannulae (Levitan et al., 2019). The presence in our dataset of neurons responding to both tastants and water could represent evidence of tactile and thus a multimodal component. Water-specific responses have been reported in many brain regions of the gustatory neuraxis including the periphery (Zocchi et al., 2017) as well as hindbrain (Rosen et al., 2010; Nakamura and Norgren, 1991) and forebrain (Gutierrez et al., 2010; Chen et al., 2021; Bouaichi and Vincis, 2020; Verhagen et al., 2003) areas. Often these water responses are classified as tactile (i.e., somatosensory) and discounted as not “taste mediated”, but the data presented in (Bouaichi and Vincis, 2020; Zocchi et al., 2017; Rosen et al., 2010), as well as our current data, argue against this view. Our analyses suggest that the spiking activity of some GC neurons contain sufficient “water-specific” information such that a pattern classifier can discriminate water from all the other stimuli. If all water responses were exclusively somatosensory (i.e., encoding common tactile inputs from all/some of the fluids within the the oral cavity), the decoding analysis would not be able to discriminate water from all the other tastes. Voltage signals from the tetrodes are acquired, digitized, and band-pass filtered (300-6000 HZ) with the Plexon OmniPlex system (Plexon, Dallas, TX) at a sampling rate of 40 KHz. Single units are then sorted offline using a combination of template algorithms, cluster-cutting techniques, and examinations of inter-spike interval plots using Offline Sorter (Plexon, Dallas, TX).

**Figure 1:**
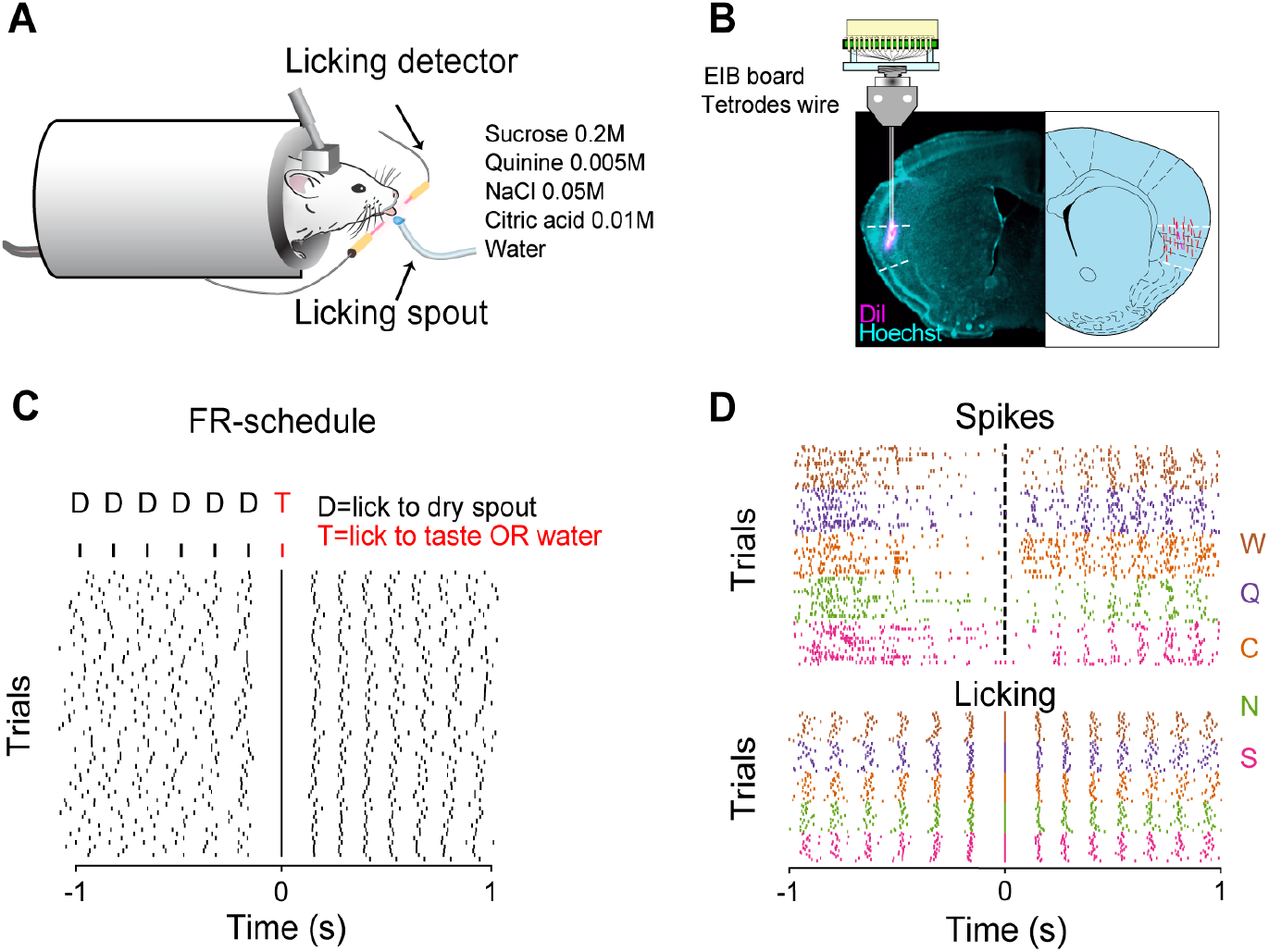
**A:** Sketch showing a head-restrained mouse licking a spout to obtain gustatory stimuli or water. **B: Left**-Example of histological section showing the tetrode tracks (magenta) in the GC; **Right**-Schematic reconstruction of the tetrode tracks of all the mice used in this study (red). In magenta the reconstruction of the track corresponding to the histological section shown on the left. **C: Top panel** Diagram of the taste delivery paradigm. Gustatory stimuli and water (T) are delivered after 6 consecutive dry licks (D) to the spout. Each lick is denoted by a vertical line. **Bottom panel** Raster plot of licking activity in a 2 s time interval centered at the taste delivery (time 0) from one experimental session. **D: Top panel** Raster plot of spiking activity from a GC neuron centered at the taste delivery, where each tick mark represents a spike. **Bottom panel** Corresponding raster plot of licking activity. Spike and lick tick marks are grouped together and color-coded with sucrose (S) in pink, NaCl (N) in green, citric acid (C) in orange, quinine (Q) in purple, and water (W) in brown.

### Experimental Design and Statistical Analysis

#### Preprocessing by binning or smoothing

Raw neuronal spike trains are represented mathematically as 4000-dimensional vectors, with one entry per millisecond of experimental data collection. Each spike train vector’s entries are valued in {0,1}, with a 1 indicating a neuronal spike at the associated time. To improve the robustness of comparisons between spike train vectors and to overcome the stochasticity that is inherent in these data, spike train vectors can be processed using *binning* or *smoothing*. Binning is a standard technique in which time is partitioned into bins, and the number of spikes per bin is determined and used as a coarse-grained representation of the spike train. An alternative approach is to use smoothing. The smoothing operation consists of replacing the raw {0,1}-valued spike train vector with a vector taking continuous positive values, achieved by convolving the raw signal with a Gaussian kernel. Intuitively, each spike in the spike train is replaced by a smooth “bump” and the spike train is represented overall as the aggregation of these bumps. This leads to a parametric spike train model, with the parameter describing the size, or the smoothing window, of each bump, with a larger smoothing window leading to a smoother functional representation. Figure 2 shows a raw spike train, together with both binned and smoothed versions of the same sample.

**Figure 2:**
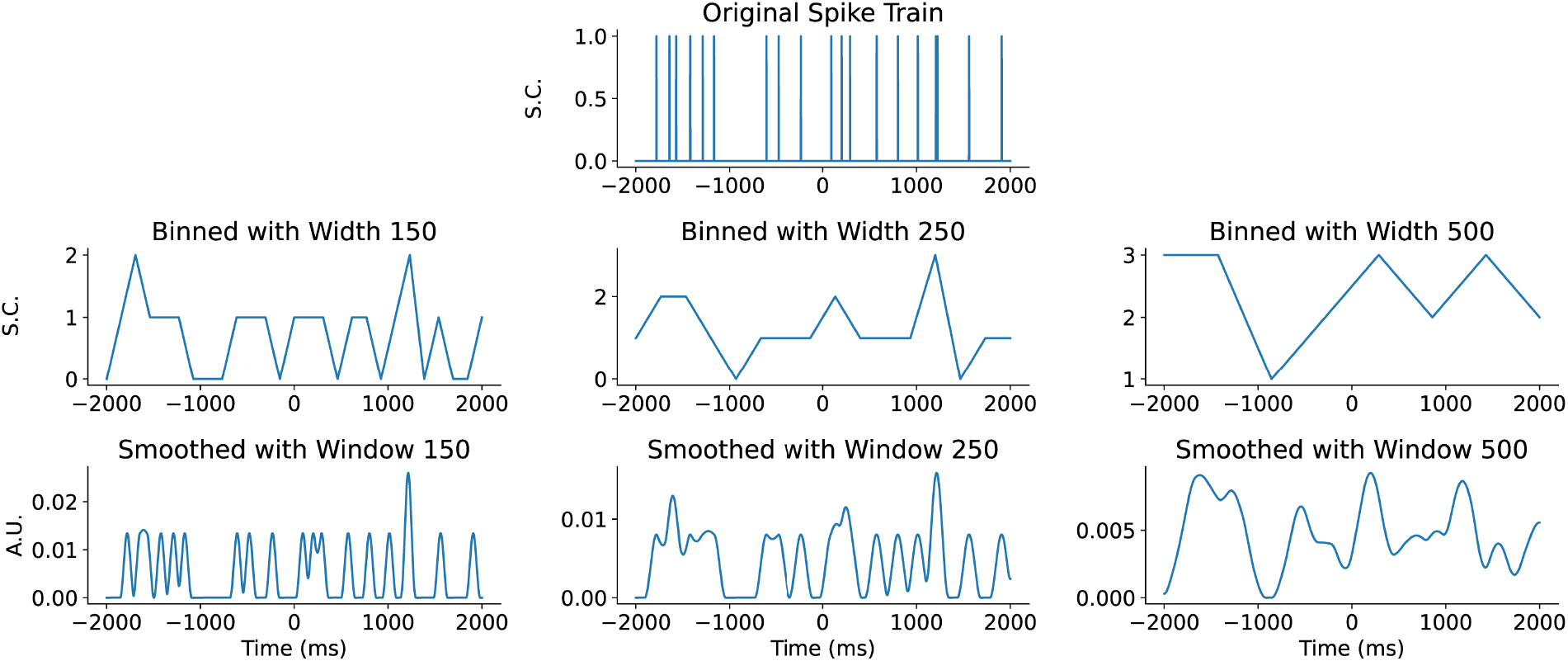
Binning and smoothing of neuronal spike trains. **Top Row:** Raw spike train consisting of a {0,1}-valued time series over 4000 ms with one spike count (S.C.) for each ms. Taste delivery occurs at time 0. **Middle Row:** Binned versions of the same raw spike train with various bin widths. Responses are a spike count for the bin. **Bottom Row:** Smoothed versions of the same raw spike train with various smoothing window sizes. Responses are in arbitrary units (A.U.).

#### Support vector machine classification

After choosing a smoothing (or binning) window, the neural spike train data set is transformed into a collection of vectors, each corresponding to one experimental trial. Each trial in the data set is classified hierarchically according to neuron ID and tastant for the trial. Thus, several vectors are associated with a single neuron, depending on how many trials were run with different tastants. To quantify single-neuron tastant decoding performance, we adopt a classical tool from machine learning called a Support Vector Machine (SVM) in order to determine a *classification score* for each neuron. Each neuron gets a classification score for the full collection of five tastants, as well as a classification score for each pair of tastants.

A classification experiment for a single neuron’s ensemble of spike trains consists of separating the ensemble of vectors into a *training set* consisting of 67% of the spike trains and a *testing set* consisting of the remaining 33% of the spike trains. The training set is used to fit parameters for the SVM model. Intuitively, the SVM searches for hyperplanes that best separate the various *classes* (i.e., points corresponding to trials for different tastants) within the data set. The trained model can then be used to classify the testing data set, resulting in a classification score for the experiment, measured as the percentage of correctly classified points. This procedure is repeated 20 times for each neuron, each time using a different partition of vectors into training and testing sets, and the classification scores are averaged over these trials to obtain an overall classification score for the neuron.

#### Rate and phase codes

To ascertain the roles of firing rate and temporal phase in single neuron tastant coding, we first introduce a simple method to reduce a spike train to a *Rate-Phase code* (RP code), consisting of two numbers. Recall that, for each trial, times at which the animal licks the tastant spout are recorded. The time axis for each neuronal spike train can then be subdivided into *interlick intervals* between these licks. The RP codes for the spike trains are constructed as follows. For all spike trains, a fixed number of 5 interlick intervals in the spike train are considered. The rate code *R* for a spike train is the average number of spikes per interlick interval considered. To compute the phase code *P*, we first compute the *mean phase* of the spikes in each interlick interval: each spike in the interval is assigned a relative position in [0,1), with with 0 corresponding to a spike concurrent with the starting lick of the interval and 1 corresponding to a spike immediately prior to the terminal lick of the interval. Finally, P is the average of these individual spike phases. The RP code for the spike train is the ordered pair (R, P). If the associated lick spike train does not contain enough licks to construct the necessary 5 interlick intervals, or in the case that the neuron does not fire in any of the interlick intervals considered, then the spike train is removed from this analysis.

To each neuron and each pair of tastants, we associate three *separation scores* to quantify the distinguishing power of the neuron for those tastants in terms of pure rate code, pure phase code and a combination of rate and phase. The *rate only*, *phase only* and *combination* separation scores are computed by determining the ability of a vertical, horizontal or arbitrarily-oriented line, respectively, to separate samples from the two tastants in the RP-plane. Separating lines are illustrated in Figure 7.

#### Elastic shape analysis

While the analysis of rate and phase codes described above gives insights on the coding methodology of sensory neurons, the reduction in complexity in transforming a full 4000-dimensional spike train to a pair of numbers is extreme. Another approach is to employ the methods of Elastic Shape Analysis (ESA) (Srivastava and Klassen, 2016), which has proven successful in many applications involving big and complex data; see, for example (Klassen et al., 2004; Amor et al., 2015; Lu et al., 2014; Marron et al., 2015) for some general applications, and (Wu and Srivastava, 2011b,a, 2012, 2013) for specific applications to neuronal spike train analysis. We employ this second approach for classification of neuron activity, in addition to the Rate-Phase code, since the classification uses the full distribution of spike timing, rather than just the mean. The numerical pipeline for taste classification of spike trains using ESA that we used was built on the open source library of Dereck Tucker, which is available on github: https://github.com/jdtuck/fdasrsf_python.

A key problem in functional data analysis consists of defining and finding *optimal registrations* – correspondences between points across data. In our setting this corresponds exactly to the problem of separating spike timing and rate. Current data analysis techniques often solve for registration as a preprocessing step, and follow it with a statistical analysis that is independent of the registration metric. That is, the “phase information” is not used. In our ESA for neuron classification, we used this phase information. The registration of a data stream requires stretching and compressing of time, a process called *time warping*. Intuitively, the aim is to find the optimal time warping of a data stream so that it best aligns with another, reference data stream.

As an example, consider two smoothed spike train functions *f* and *g* defined over the time interval [0,T]. There are many ways to measure the distance between functions of this form; we choose the *extended Fisher-Rao metric* (Srivastava and Klassen, 2016; Bauer et al., 2016), defined by

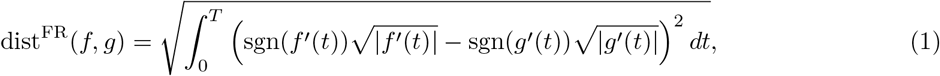

where *f*’ denotes the derivative of the function *f* and sgn is the sign function—i.e., sgn(*s*) = 1 if *s* ≥ 0 and otherwise sgn(*s*) = −1. The reason for choosing this metric is that it is invariant under time warpings of the data. A *time warping* is represented mathematically as a smooth invertible function *γ* with positive derivative over the domain [0, *T*]. The time-warping of a function *f* by the warping function *γ*, which we refer to as an *alignment function,* is given by function composition, denoted as *f* ◦ *γ*. For the Fisher-Rao metric, *invariance under time warpings* means that

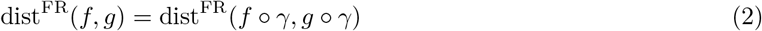

for any alignment function *γ*. We consider a signal *f* to have the same *rate code* as any time warping *f* ◦ *γ*. The distance between the rate codes of signals *f* and *g* can be computed by solving the optimization problem

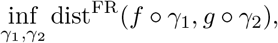

where the infimum (i.e., minimum) is over pairs of alignment functions. Due to the invariance property (2), this is equivalent to minimizing the distance dist^FR^(*f,g* ◦ *γ*) over a single alignment function *γ*. In practice, this is solved by discretizing the time domain and the input signals *f* and *g* and optimizing over *γ* using a dynamic programming algorithm. For the minimizing alignment function *γ*, the traces of the signals *f* and *g* ◦ *γ* give a visualization of the similarities and differences between the rate codes of the signals *f* and *g*, whereas the trace of *γ* illustrates the relative phase differences between the signals.

Figure 3 shows two examples of optimal alignments between signals (smoothed spike trains) using ESA. In the first example, the two signals have the same rate, but differ significantly in phase (panel A). After alignment they are identical (panel C). Thus, the alignment yields two signals with the same number of peaks, indicating that the spike rates are identical, and the alignment function (panel B) applied to the blue trace indicates how the timings differed from those of the red trace. The second example shows a case where the smoothed signals have different rates. In one case (panel D, red), spikes are clustered together so that the signal after smoothing is a single large peak. In the other, the same number of spikes are separated sufficiently so that there are two distinct peaks in the smoothed signal (panel D, blue). The smoothed signals are therefore different in both rate and timing. The alignment function (panel E) applied to the blue trace cannot completely reconcile the two signals; the red signal has a single peak while the aligned blue signal still has two peaks, but is now aligned with the larger red peak. The differences in spike rate of the smoothed signals is preserved.

**Figure 3:**
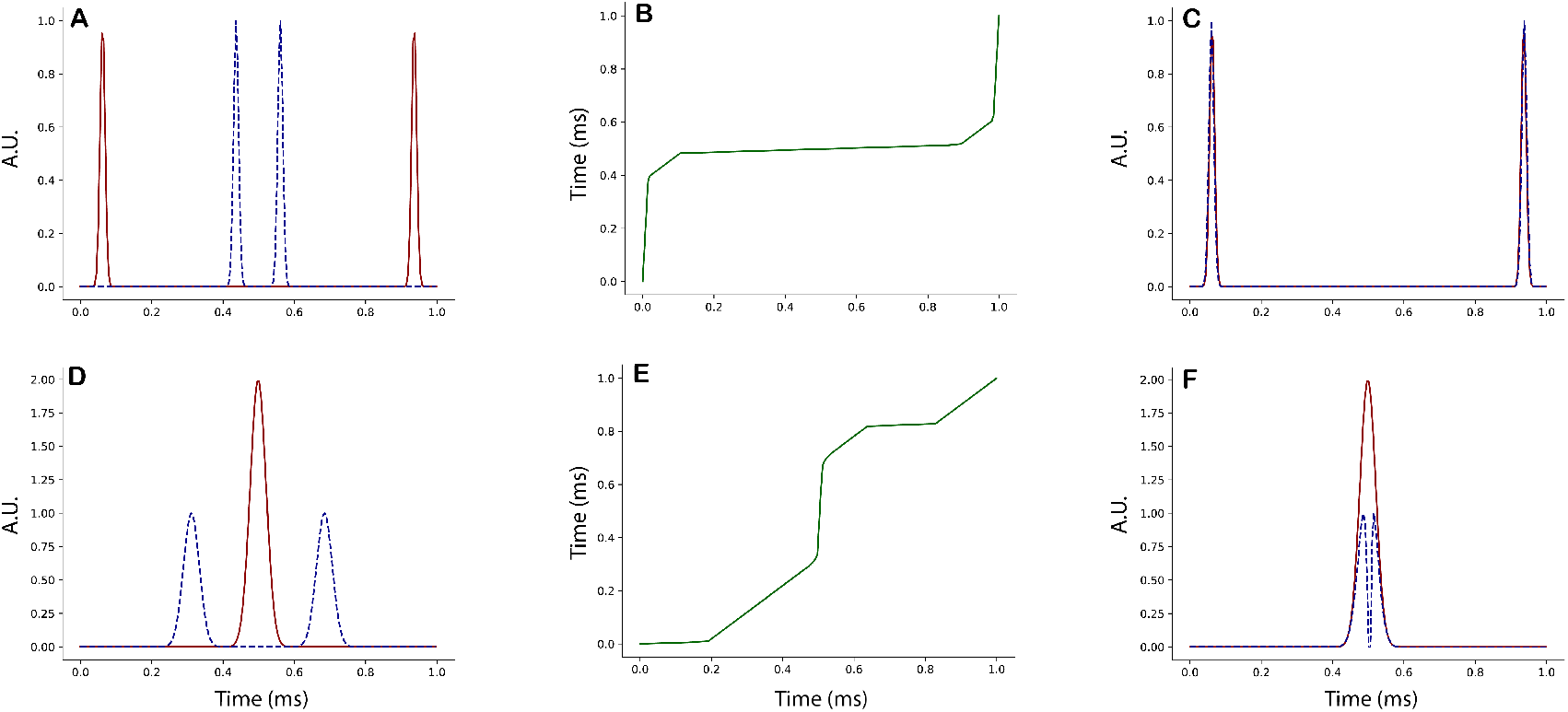
Illustration of the behavior of the Fisher-Rao metric. Each row shows a different example of aligning a pair of synthetic signals (i.e., solving the optimization problem described in the text), with responses measured in arbitrary units (A.U.). Panel **A** shows a pair of signals *f* (red, solid) and *g* (blue, dashed). Considering these signals as smoothed spike trains, it is clear that the spike trains have the same firing rate, but that they differ in phase. This is made precise using the ESA framework. Solving the ESA optimization problem produces an optimal alignment function *γ*, as shown in panel **B**. The time warped signal *g* ◦ *γ* is shown in panel **C** (blue, dashed), together with the signal *f* (red, solid)—the traces are the same, indicating that f and *g* have the same rate code. The fact that the optimal alignment function *γ* is far from the identity tells us that the signals *f* and *g* differ significantly in phase. Panel **D** shows another pair of functions *f* (red, solid) and *g* (blue, dashed), and the optimal alignment function *γ* is shown in panel **E**. Observe that the graphs of *f* and *g* both enclose the same area; however, *f* has a single maximum, whereas *g* has two local maxima. Following alignment, there are still two peaks in *g* ◦ *γ* (panel **F**), versus one in f, so the difference in rate is preserved.

After the registration process we have represented our data as *f* and *g* ◦ *γ*, and the corresponding alignment function *γ*. To use this method in combination with SVM classification, we align all spike trains to the *mean spike train* of the population (i.e., a function that is the centroid of all spike trains, minimizing the sum of squared distances to the samples with respect to the extended Fisher-Rao distance (1)). An SVM classification is then performed on these aligned smoothed spike trains to determine performance using **rate only coding**, since the phase component was removed by the alignment. A separate SVM classification is also performed, this time using the alignment functions themselves rather than the smoothed spike trains, providing a measure of performance using **phase only coding**.

## Results

### Using a machine learning algorithm for single-neuron taste discrimination

To investigate how neurons within the primary taste cortex encode gustatory information in freely licking mice (Fig. 1A), we made recordings using bundles of tetrodes implanted unilaterally in the GC (Fig. 1B). After habituation to head restraint, water-deprived mice were engaged in a task in which they had to lick a dry spout 6 times to obtain a 3 μL drop of one of five gustatory stimuli (sucrose, 200 mM; NaCl, 50 mM; citric acid, 10 mM; quinine, 0.5 mM; water) (Fig. 1C). Mice were trained until the licking pattern evoked by each of the individual stimuli was similar across a 1 s temporal epoch following taste delivery (Fig. 1D, see (Bouaichi and Vincis, 2020) for more details). To begin exploring the neural dynamics evoked by intra-oral stimuli, we first evaluated using Support Vector Machines (SVM) as a tool for decoding taste information from single neurons. This process was run using trials from all 5 tastants and using trials from 2 tastants (running over all possible pairs).

#### Comparison of binning and smoothing for SVM classification

The calculation of the classification score is influenced by how the spike train data are represented. The raw form, essentially a collection of action potential times, has very little overlap of spike trains regardless of the tastant applied, so while it provides sufficient information for rate-code classification, it is not optimal for classification that uses temporal overlap of neural activity as well as rate. A more appropriate way to treat the data is to use binning. The spike train is thereby processed into a vector where each element contains the number of spikes in a bin. The bin width (in ms) is a free parameter that determines the size of the vector used to represent the spike train, and will affect the classification score. This is demonstrated in Fig. 4, using 10 different bin sizes ranging from 1 to 500 ms (where a size of 1 ms means that the unbinned spike trains are used). The figure shows the classification score using all 5 tastants. When spike trains from the 10% of best-performing neurons are analyzed, the classification score improves from ~0.25 to ~0.4 when the bin window is increased. A greater improvement is seen when only the top 5% performing neurons are used, and when the top 1% are used the classification score rises from ~0.32 to ~0.55.

**Figure 4:**
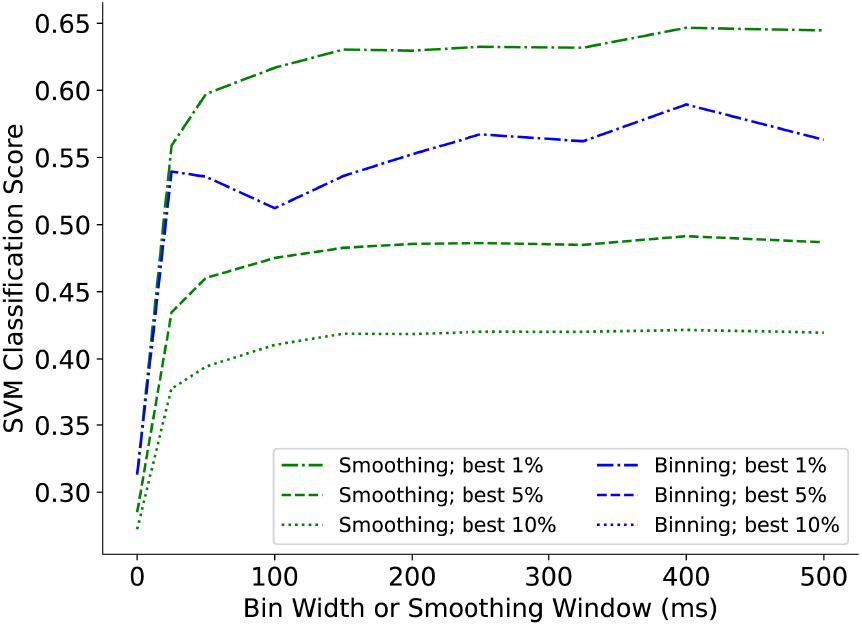
SVM classification scores of all 5 tastants and full length spike trains treated with varying smoothing and binning parameters. At each parameter level, the mean classification rate among the best-performing 1%, 5%, and 10% of neurons is plotted.

An alternate approach for processing the spike train data is to smooth it by convolving a spike train with a Gaussian function, converting the spike times into a series of bumps that can accumulate when they occur close together in time. When this approach is used, the classification score increases as the smoothing window is increased (Fig. 4), similar to the case of binning. However, the improvement in classification score is even greater with smoothing, and for all window values used the classification score is higher using smoothed data than using binned data. For example, when the top 1% of the neurons are used, the classification score is ~0.65 for smoothed spike trains vs. ~0.55 for binned spike trains. An additional feature of smoothing is that the classification score is stable for all window values of 100 or greater, so the classification results do not depend strongly on the smoothing parameter value, as long as it is sufficiently large.

#### SVM with all five stimuli

Based on the results above (smoothing vs. binning), we used smoothed spike trains with smoothing window of 250 ms to represent our data. We first investigated the distribution of SVM classification scores for 5 stimuli across the population of GC neurons. For each neuron, all of the smoothed spike trains were used to compute a classification score. A classification score of 1 means that all trials in the testing set for a neuron were properly classified according to the tastant. The classification scores for all neurons are shown in the top row of Fig. 5; each neuron has an SVM classification score based only on its pre-taste signal, and another based only on its post-taste signal. In panel A, a histogram shows the distributions of pre-taste classification scores (in yellow) and post-taste scores (in pink and blue). Panel B focuses only on the post-taste scores.

**Figure 5:**
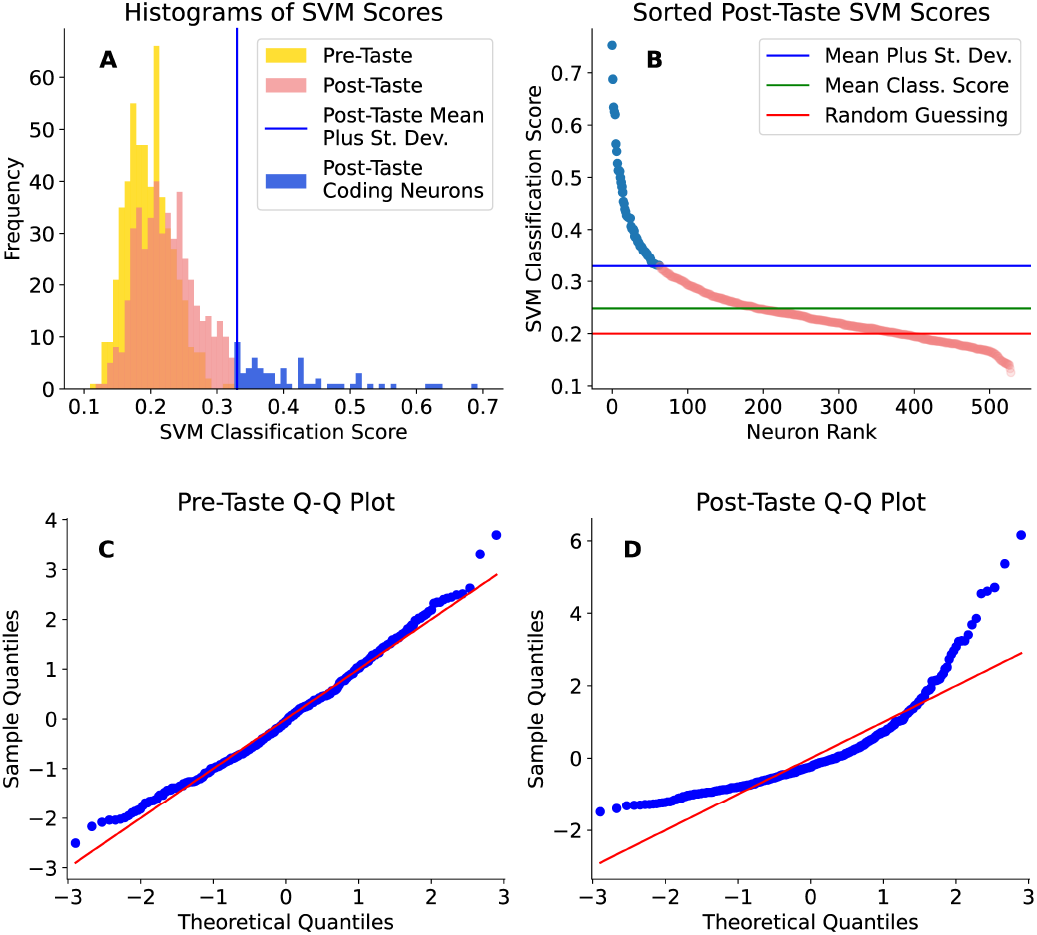
**Top Row:** Histogram of the SVM classification scores for all 5 tastes on the pre-taste and post-taste data (**A**) and a closer look at the post-taste scores (**B**). High post-taste scores are colored in blue, and the corresponding neurons are referred to as “coding neurons”. **Bottom Row:** Quantile-Quantile plots for the SVM scores using pre-taste data (**C**) and post-taste data (**D**). For reference, the red lines indicate the quantiles of a normal distribution.

To analyze the distributions of classification scores, we used two statistical tests: D’Agostino and Pearson’s normality test (D’Agostino, 1971; D’Agostino and Pearson, 1973) and a one-sided t-test on the mean classification score. Both tests were run separately on the pre-taste and post-taste scores. On the pre-taste set, D’Agostino and Pearson’s normality test showed the classification scores are not normally distributed (*K*^2^ statistic=12.278, p-value=0.00216 for *H*_0_: the sample is normally distributed). However, (*K*^2^ statistic = 12.278, p-value = 0.00216 for *H*_0_: the examination of the quantile-quantile plot in Fig. 5C shows only a slight departure from the theoretical normal quantiles occurring mostly at the tails. A departure from normality in the sample distribution does not, however, render a t-test invalid. Validity of this test requires that the sample means are normally distributed, not the sample itself. This condition is met for sufficiently large samples due to an application of the Central Limit Theorem (Kwak and Kim, 2017). In addition, t-tests are relatively robust to deviations from this normality assumption (Sawilowsky and Blair, 1992). When a t-test was performed, the mean successful classification of pre-taste spike trains was determined to be no better than random guessing (t-statistic=-1.76, p-value=0.9602 for *H*_0_: *μ_pre_* = .2 and *H_A_*: *μ_pre_* > .2).

In contrast to the pre-taste scores, the classification scores for post-taste spike trains show a much greater skewness towards good-performing neurons (Fig. 5B). D’Agostino and Pearson’s test on the post-taste classification rates show a very large departure from normality (*K*^2^ statistic=265.443, p-value=2.290× 10^-58^ for *H*_0_: the sample is normally distributed). Examination of the quantile-quantile plot (Fig. 5D) shows a large departure from the theoretical quantiles across the entire distribution. The mean of these classification rates is significantly above random guessing (t-statistic=13.428, p-value=7.08×10^-36^ for *H*_0_: *μ_post_* = .2 and HA: *μ_post_* > .2). These results demonstrate that the neural signals before taste administration are little more than noise, but that there is indeed a significant signal in the post-taste data. In particular, there are approximately 50 (or 10% of the neurons analyzed) that are “coding neurons” in the sense that they successfully classify information from all five tastants, demonstrated by a score more than one standard deviation greater than the population mean.

#### SVM with taste pairs

We next investigated a binary classification task, testing the ability of the SVM applied to neural spike trains to distinguish between a pair of taste stimuli. In this test, training of the SVM was performed on spike trains in response to each of two tastants and SVM classification scores were obtained for each combination of neuron and taste pair. The results for selected taste pairs are shown in Fig. 6 (A-D). The key findings are similar to those from the analysis with all tastants: while the pre-taste classification scores were mostly noise, the post-taste classification scores were skewed and had a mean that is significantly above random guessing. For spike trains of some neurons, the SVM was not successful at distinguishing between two tastes. However, for many of those neurons in which the SVM was successful (the top 10-20%), the classification score was very good. There is also a significant amount of overlap between the coding neurons for different taste pairs. As shown in the cumulative distribution histogram of Fig. 6E, approximately 60% of the total population of neurons studied are coding neurons for at least one taste pair (i.e., the classification success rate is greater than one standard deviation from the mean). Approximately 30% of the neurons are coding neurons for at least two taste pairs, and approximately 20% are coding neurons for at least three taste pairs. The very best neurons (1% of the population) are coding neurons for all 10 taste pairs.

**Figure 6:**
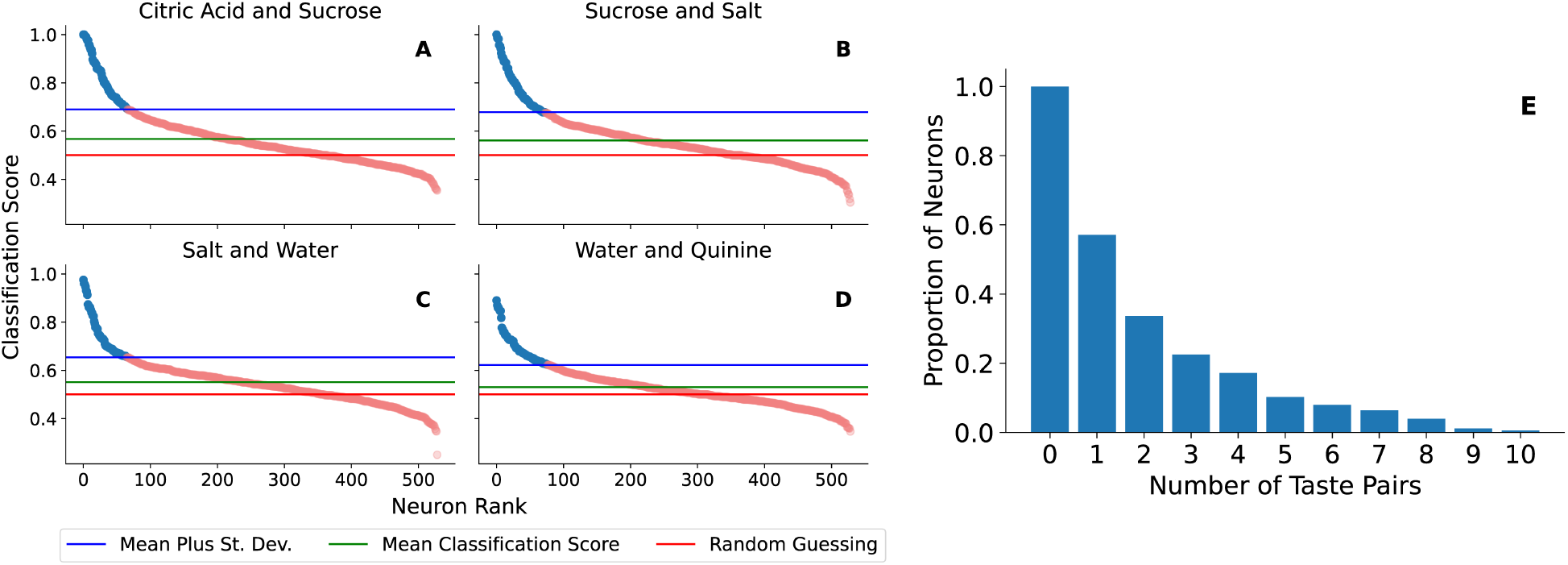
**A-D:** Support Vector Machine classification rates on selected taste pairs using post-taste test set spike trains. The distributions are heavily skewed towards good-performing coding neurons, colored in blue. **E:** A cumulative distribution histogram of the fraction of neurons that are labeled as coding neurons for one or more taste pairs.

Overall, these first set of results indicate that SVM classifiers are an effective method for quantifying taste information and for identifying neurons that best encode that information.

### Using averaging to determine the contributions of spike rate and phase in taste classification

The next task is to determine the relative contributions of spike rate and phase to overall decoding performance. We begin by using a Rate-Phase code (RP code) that is based on the mean spike rate (R) between consecutive licks and the mean phase (P) at which spiking occurs between consecutive licks, as discussed in Methods.

Each spike train is divided into interlick intervals [*t*_*i*–1_, *t_i_*), *i* = 1,..., 5, where *t*_0_ = 0 is the time of the first lick at which a tastant was administered and *t_i_* denotes the time of the ith subsequent lick. The rate code *R* is then the average number of spikes per lick interval. If a spike occurs at time *t* in interlick interval [*t*_*i*–1_, *t_i_*), the phase for the spike is given by (*t* – *t*_*i*–1_)/(*t_i_* – *t*_*i*–1_). The phase code P for the full spike train is the average of these individual phases over all spikes.

Once the RP codes were computed for a neuron, information for each spike train was plotted as a point in the RP-plane. This resulted in a cloud of points in the RP-plane, reflecting the neuron’s responses to two different tastants as demonstrated in Fig. 7. We then searched for the line that separated the greatest number of observations based on taste, using three methodologies. Scanning over vertical lines, we obtained a *rate separation score*: for each vertical line, we compute the maximum of

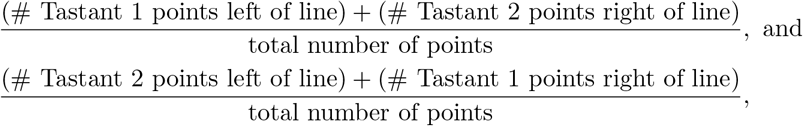

and defined the rate separation score as the maximum possible value over all vertical lines. This quantifies how well the neuron is able to separate the tastes using only average interlick firing rate information. We computed a *phase separation score* by scanning across horizontal lines and performing a maximization using a similar formula. This quantifies the ability of the neuron to distinguish the two tastes using only average interlick phase information. Finally, we computed a *combination separation score* by scanning over *all* lines of arbitrary slope and maximizing a similar function, thereby quantifying the ability of the neuron to distinguish the tastes using both average interlick firing rate and phase information. By construction, the combination score will be at least as good as the rate or phase score alone, cf. Fig. 7.

**Figure 7:**
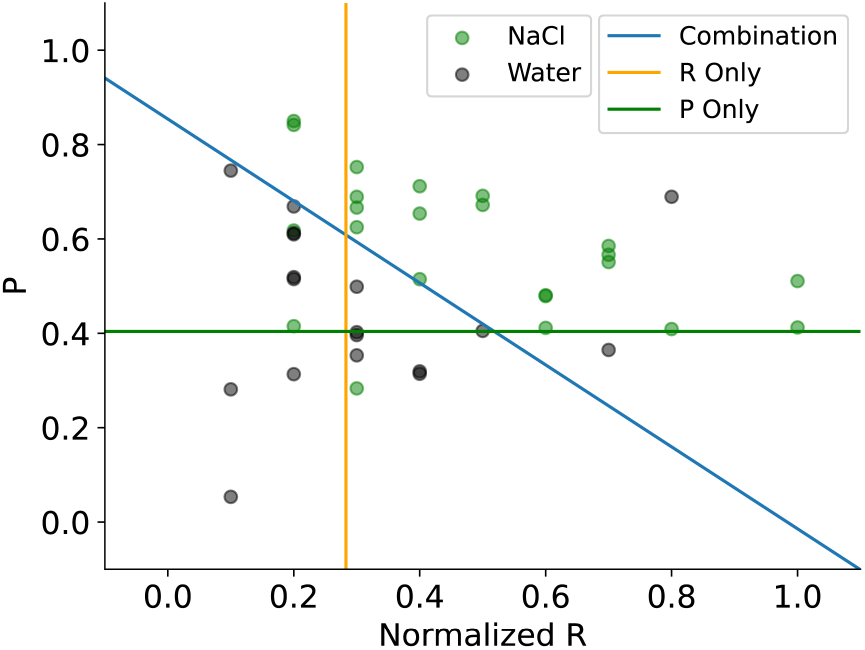
Rate-Phase code for the two tastants Water and NaCl. The orange line shows the best separation of the data based on mean rate only, the green line shows the best separation based on mean phase only, and the blue line is the best separating line based on both mean rate (R) and mean phase (P).

Figure 8 shows summary results for four taste pairs. In each, separation scores for the 50 neurons with the best combination separation scores are shown. For all of these, 80% or more of the points could be successfully separated using the simple RP code. Thus, the combination of mean spike rate and phase was quite successful at clustering spike trains according to the taste stimulus. For some neurons and taste pairs, the best separation was achieved using pure rate separation (orange dots) or pure phase separation (green dots), but in the vast majority of cases the best separation required a combination of the two (blue dots). Among the better neurons, the rate separation score was better than the phase separation score: among the top ten neurons the average rate separation score was better than the phase separation score.

**Figure 8:**
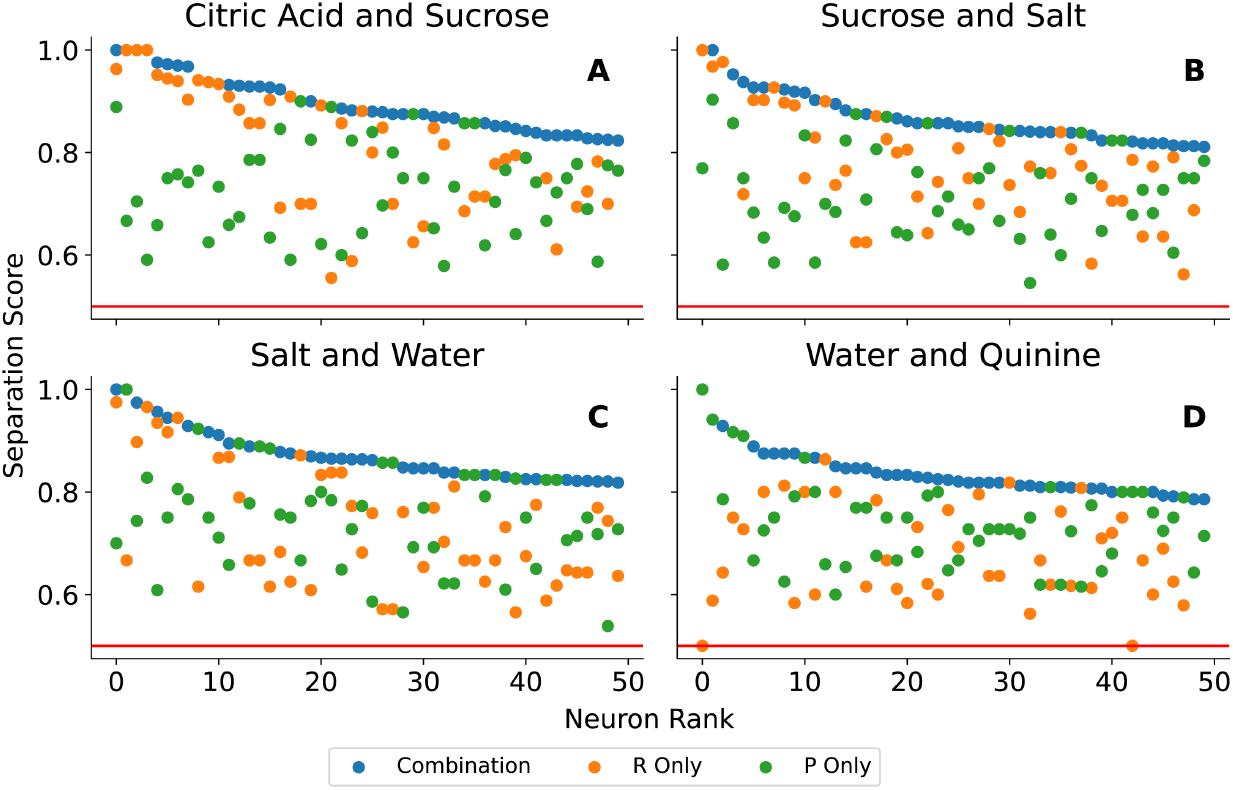
Separation scores for selected taste pairs applied to the best 50 neurons for each taste pair.

The separation scores using either R or P alone become similar after the first ~ 25 neurons. In fact, among the top 50 neurons for each taste pair on average the rate separation score is only better in 23.5 cases. The separation scores for rate decrease quickly over the best neurons, while the scores for phase stay more constant. To quantify this, we calculated the mean R and P-score for the best 10 and 50 neurons for all 10 taste combinations: for the 10 best neurons the mean R-score is in six cases (Citric Acic–Sucrose, Citric Acid– Salt, Citrid Acid–Water, Sucrose– Quinine, Salt–Quinine and Sucrose–Salt) significantly better (p-value in the t-test below 0.05). Repeating the same analysis for the best 50 neurons we obtain that the mean R-score is only in three cases (Citric Acic–Sucrose, Citric Acid–Salt and Sucrose–Salt) significantly better (p-value in the t-test below 0.05), while the P-score is in one case significantly better (Water–Quinine). For the remaining cases no statistically significant statement can be made.

### Using elastic shape analysis to quantify rate and temporal/phase coding

The simple RP code used in the previous section employs an extreme dimension reduction of the data that provides an intuitive metric for quantifying the contributions of rate and phase coding to taste discrimination. However, the dimension reduction averages out many features of spike train timing. In this section we employ a functional data analysis technique that does not use this averaging step, and thus is more appropriate to properly assess spike timing information. In particular, we use the Elastic Shape Analysis (ESA) technique to find the optimal alignment of a group of spike trains, as described in Methods. To evaluate the relative contribution of rate and temporal/phase coding in GC neurons, we perform ESA alignment in two ways: with no consideration of when licks occur and by using the licks to partition time into interlick intervals. Since the latter alignment considers spike timing relative to the licking times, it provides phase information.

#### ESA with unconstrained alignment

With the first application of ESA, disregarding lick timing, an “average spike train” is computed for each neuron, where the averaging is done over the set of all post-taste spike trains for that neuron (specifically, the average is a function that minimizes the sum of squared distances to the samples, with respect to the Fisher-Rao metric). Then, each spike train is optimally aligned to the average spike train. The optimization is performed by determining an alignment function, as described in Methods. Thus, for each neuron, the ESA process results in a collection of aligned post-taste spike trains and an accompanying collection of alignment functions. The aligned spike trains lack accurate spike timing information (it was removed by the alignment), but contain rate information. The alignment functions contain information on when spikes occurred relative to those in the mean spike train, and thus contain spike timing information. It is therefore possible to perform an SVM classification using the aligned spike trains as well as a second SVM classification using the alignment functions themselves, thereby determining classification scores using spike rate and spike timing, respectively. The terminology that we use for the spike trains and alignment functions is summarized as:

- **Original (smoothed) spike train:** the spike train smoothed with a smoothing window of 250 ms. Contains rate and spike timing information.
- **Aligned spike train:** contains rate information for the 2 s long post-taste interval without regard to lick timing
- **Alignment function:** contains spike timing information for the 2 s long post-taste interval without regard to lick timing
- **Interlick aligned spike train:** contains spike rate information within lick intervals
- **Interlick alignment function:** contains spike timing information within interlick intervals (i.e., phase information)

Figure 9 shows examples in which this ESA technique is used to classify spike trains in response to four taste pairs. In each panel, the 50 best neurons determined using the SVM classification scores for the original spike trains (which contain both spike rate and spike timing information) are shown in blue along with classification scores using the aligned spike trains (orange) and the alignment functions (green). For many neurons, the classification score using the aligned spike trains is almost as good as that using the original spike trains, and in a few cases even better. For example, in the case of citric acid versus sucrose, the classification score for the aligned spike trains is at least as high as that for the original spikes train in 18 out of 50 neurons, and on average the aligned spike train score is 96.2% as large as the original spike train score. The classification scores based on the alignment functions are somewhat lower. In the case of citric acid versus sucrose, this score is greater than that using the original spike trains in 4 out of 50 neurons and on average is 79.5% as large as the classification score using the original spike trains. These analyses show that using both spike rate and spike timing gives better classification for taste pairs. When the aligned spike trains (which preserve spike rate) are used in the classification, the classification score goes down, but is better than when only the alignment functions (which provide the spike timing information) are used in the classification.

**Figure 9:**
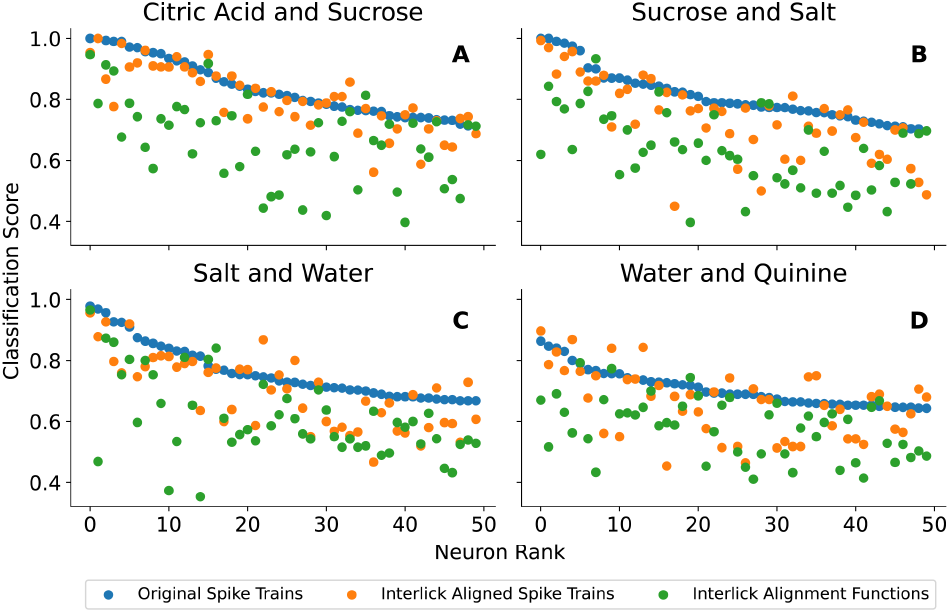
SVM classifiction on four selected taste pairs. In each case, the 50 best neurons based on SVM classification of original spike trains are used and the classification score is in blue. The classification scores using the aligned spike trains are in orange, and those using the alignment functions are in green. The ESA and SVM classification are performed on spike trains after administration of the tastants and without consideration of lick timing.

#### ESA with alignment performed on interlick intervals

For the second ESA experiment, we performed the alignment on each interlick time interval. Since we use the first 5 post-taste interlick intervals in the analysis, each spike train is split into 5 aligned spike trains and 5 alignment functions. These are then concatenated to form a single post-taste aligned spike train and a single interlick alignment function. The interlick alignment functions are now an indicator of spike phase relative to licks, so using SVM classification on these alignment functions demonstrates how well tastes are differentiated using a pure phase code. The results using four taste pairs are shown in Fig. 10, which is formatted in the same way as Fig. 9.

**Figure 10:**
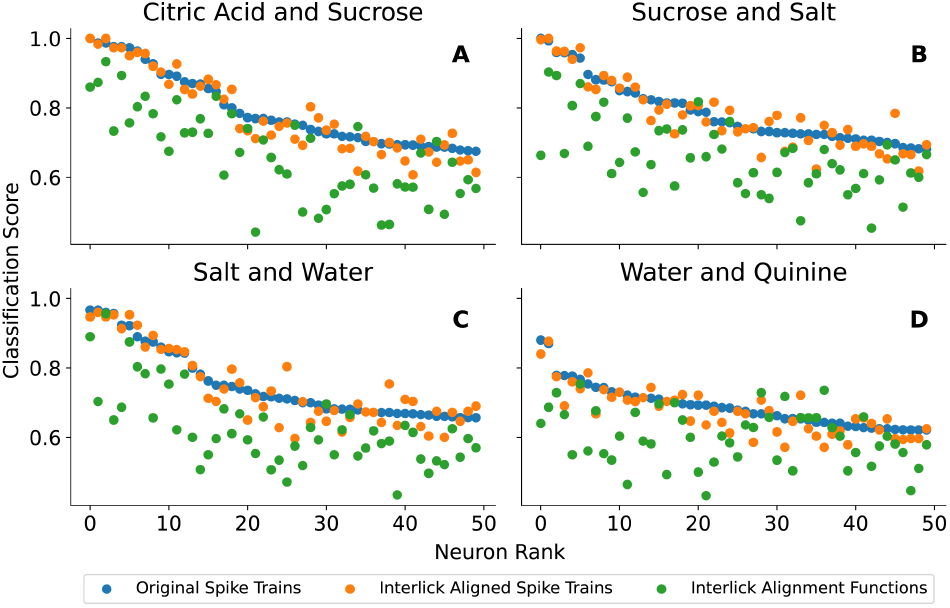
SVM classification on four selected taste pairs in which alignment is performed over interlick intervals. In each case, the 50 best neurons based on SVM classification of original spike trains are used and the classification score is in blue. The classification scores using the interlick aligned spike trains are in orange, and those using the interlick alignment functions are in green.

Once again, SVM classification using the original spike trains typically performed best (blue circles), and classification using the interlick aligned spike trains (orange) typically outperformed classification using the interlick alignment functions (green). For example, in the case of citric acid versus sucrose, the classification, score for the interlick aligned spike trains is at least as high as that for the original spike train in 19 out of 50 neurons, and on average the interlick aligned spike train is 98.5% as large as the original spike train. For classification using the interlick alignment functions, the score is greater than that using the original spike trains in 2 out of 50 neurons and on average is 83.8% as large as the classification score using the original spike trains.

#### ESA on all five stimuli

Next, we analyzed the classification ability for all 5 tastants, rather than for tastant pairs (Fig. 11). In this case, random guessing would result in an average classification score of 0.2. For the 50 best neurons (based on the original spike trains), the classification was always much better than what would be expected from random guessing.

**Figure 11:**
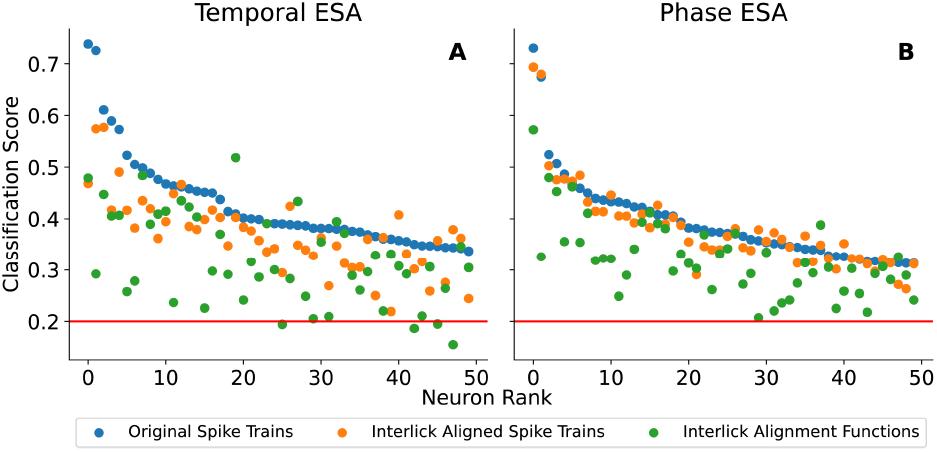
SVM classification scores for all 5 tas-tants. The classification uses either original spike trains (blue), aligned spike trains (orange), or alignment functions (green). **A:** Alignment is performed over the entire 2 s post-taste time interval. **B:** Alignment is performed on five individual post-taste interlick intervals.

As in the two previous figures, the best classification scores were typically obtained when the original spike trains were used in the classification (blue) followed by classification scores with aligned spike trains (orange) and finally classification using the alignment functions (green). In panel A, the alignment was applied over the entire post-taste time period, without regard to lick timing. In panel B, it was applied separately to each interlick interval. Both the classification scores using aligned spike trains and using alignment functions were generally closer to those using the original spike trains in the latter case, where the alignment was applied to interlick intervals. On average, the classification score for the aligned spike train is 87.3% as large as that for the original spike train when aligning over the entire post-taste time interval, while it is 97.0% as large when aligning over interlick intervals. Similarly, the average classification score using the alignment functions relative to that using the original spike trains was 75.9% as large when alignment was performed over the entire post-taste time interval, and a higher 82.8% when performed over individual interlick intervals (and thus providing spike phase information).

#### ESA constrained to random intervals

The data in Fig. 11 show that the SVM classification scores for aligned spike trains and alignment functions do not drop when the post-taste data were partitioned into 5 interlick intervals. This could be explained in either of two ways: 1) irrespective of the lick partitioning, in the phase ESA the taste information is still high because we are analyzing a 5-lick post taste temporal window that could contain more chemosensory-related taste information (Katz et al., 2001) (as opposed to the “full” 2-s long post stimulus interval in the Temporal ESA) or 2) the spike rate and spike timing relative to the lick interval contained non-trivial taste information. To determine which explanation is true, we next ask whether a similar taste decoding performance is achieved by splitting the 5-lick-long post taste temporal window spike trains into 5 randomly chosen time intervals (112.46±20.47 ms). That is, we still split up the 5-lick post taste interval into 5 sub-intervals, but without regard to when licks occur. The randomized partitioning was performed 20 times for each neuron, and the resulting classification scores of these 20 experiments were averaged to mitigate the effects of the particular random partitions chosen. As shown in Fig. 12A, the classification scores are lower when the spike train is partitioned into five random intervals than when the neural activity is partitioned into five lick intervals (paired t-test; see Table 1). Thus, taste decoding in terms of either rate or phase is more effective when the time intervals on which ESA is performed correspond to the interlick intervals. Overall, our results suggest that the lick cycle is a key factor for taste processing.

**Figure 12:**
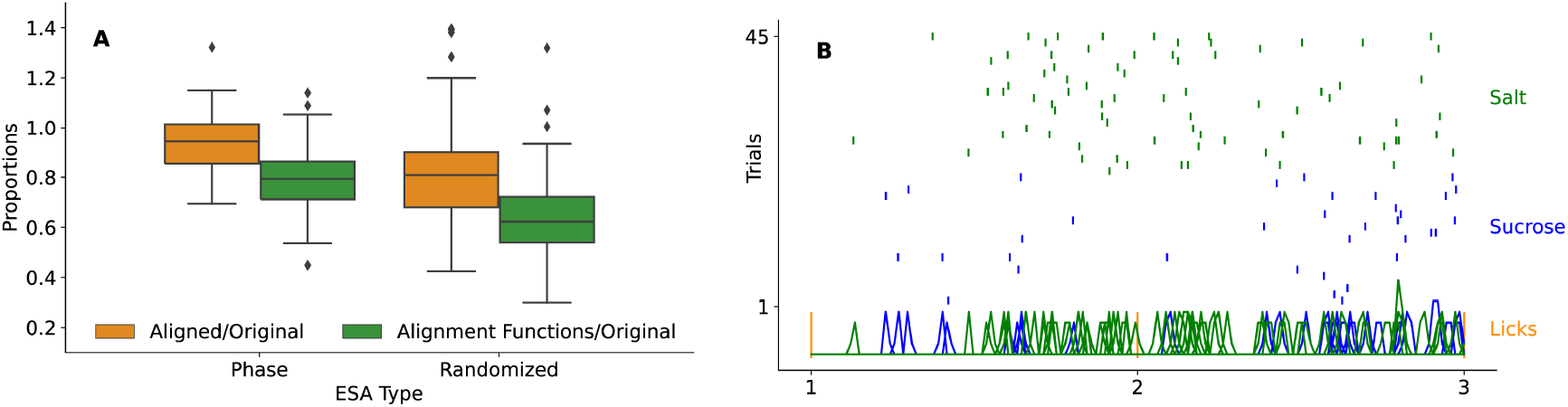
**A:** Box plot depicting the proportion of the “rate” (aligned; orange) and “temporal” (alignment functions, green) scores to the original SVM score, fromtwo different types of ESA experiments (phase and randomized). Left boxes boxes (phase): ESA is performed on spike trains in 5 interlick intervals;, right boxes (randomized): ESA is performed on spike trains in 5 random intervals. Table 1 below provides the information of each statistical comparison. **B:** Example raster plot of spiking during the first two interlick intervals in response to two tastants. At the bottom, the spike timings over all trials are represented as smoothed green or blue bumps. The overall SVM classification score for this neuron was 90%, and when classification was performed using aligned spike trains in interlick intervals the score was 96.3%, and when using the alignment functions in the classification the score was 89.3%.

**Table 1:** The statistics and p-values associated with the comparison between the ESA types in Fig. 13A. The neurons shown in the Phase ESA plot were used in the Randomized experiment, so there is a direct neuron-to-neuron comparison. *μ_D_* denotes the mean of the differences between each neuron’s Phase ESA Score and Randomized ESA Score. The appropriate test to compare the samples was a paired t-test with *df* = 49, *H*_0_: *μ_D_* = 0, and *H_A_*: *μ_D_* =0.

| ESA Types | Score Type | t-statistic | p-value |
| --- | --- | --- | --- |
| Phase vs. Randomized | Aligned/Original | 8.7596 | $1.3490 \times 10^{-11}$ |
| | Alignment Functions/Original | 8.9008 | $8.2896 \times 10^{-12}$ |

## Discussion

Multiple studies have been performed over the past three decades on the taste response profile of GC neurons in awake rodents (Samuelsen and Vincis, 2021). Extracellular recordings from animals receiving taste stimuli administered through intraoral cannulae highlighted the importance of seconds-long temporal dynamics of taste-evoked spiking activity (Katz et al., 2001). Neural data recorded from rodents that actively lick a spout to receive taste stimuli revealed further rich and interesting temporal dynamics (Bouaichi and Vincis, 2020; Gutierrez et al., 2010; Stapleton et al., 2006). One observation was that cortical taste responses displayed shorter latency when the stimulus was obtained through licking, suggesting that the temporal dynamics may vary depending on whether a taste is obtained passively or actively. Another observation was that the spiking activity of a large proportion of GC neurons display lick-coherent activity (spiking in the 5-10 Hz frequency domain) (Bouaichi and Vincis, 2020; Stapleton et al., 2006), suggesting that the stereotypical rodent licking activity may act as a metronome against which GC spikes can be coordinated (Gutierrez et al., 2010). Based on this latter point, we hypothesized that 1) changes in the rate of spike trains within “lick-cycle-long” time intervals (as opposed to rate changes over randomly defined and lick-unrelated temporal windows) and 2) the phase of spiking relative to lick timingare good discriminators of taste that can be used by GC neurons in active licking mice.

Consistent with previous reports (Levitan et al., 2019; Bouaichi and Vincis, 2020), our results indicate that GC neurons can discriminate between the different taste stimuli using changes in the spiking rate over a long post-taste temporal window. Our analysis tools allowed us to quantify the rate contribution separately from the temporal contribution to coding. While clearly the dominant factor, rate information was complemented by the timing of spikes in almost all identified coding neurons (Figs. 8–12).

It has long been known that neurons can encode sensory information via the number of spikes within a given time interval (Adrian, 1926), but the time intervals used are often chosen arbitrarily. In our experimental settings, mice actively sample taste stimuli via the rhythmic and stereotypical behavior of licking (Travers et al., 1997) that can serve as a metronome allowing GC neurons to extract and analyze sensory information in discrete interlick time intervals. Indeed our analysis indicated that the taste information contained in the spike count of GC neurons is higher when rate is computed over interlick intervals (Figs. 11, 12). If the information needed by GC neurons to discriminate different tastants is contained in short temporal intervals (such as between licks), it could allow the animal to rapidly discriminate between two different taste cues. This is in agreement with behavioral studies showing that rodents can discriminate different taste qualities using the information contained in one to two licks (Halpern and Tapper, 1971; Graham et al., 2014) and that licking-induced synchronicity in multiple brain areas of the taste-reward circuits plays an important role in taste-guided discrimination tasks (Gutierrez et al., 2010).

The timing of action potentials within a specific time window can provide substantial additional information of a stimulus (MacKay and McCulloch, 1952; Bair and Koch, 1996). Studies of the somatosensory and olfactory systems in active sensing rodents have indicated that the spike timing relative to whisking (Curtis and Kleinfeld, 2009) or sniffing can contribute to the neural representation of sensory stimuli (Smear et al., 2011; Shusterman et al., 2011). In the taste system, studies have shed some light on the potential role of neural activity time-locked to the lick cycle (Gutierrez et al., 2010; Roussin et al., 2012; Bouaichi and Vincis, 2020; Stapleton et al., 2006). Building on this, our analysis revealed that spike timing relative to the lick cycle (phase coding) contributes to taste discrimination in all of the GC coding neurons examined. Both the Rate-Phase analysis that employs averaging and the functional data analysis that does not (see Figs. 7 and 11) revealed that the phase contains sufficient information to distinguish among taste stimuli, providing quantitative evidence in favor of the importance of fine-scale temporal neural dynamics for taste-evoked activity. To evaluate the importance of lick timing in the neural coding, we performed separate analyses in which lick timing was employed and when it was not. We found that taste discrimination improved when the rate and timing information was extracted from interlick intervals, rather than random intervals of similar duration (Fig. 12A).

One challenge in the interpretation of our data is that, consistent with other studies in behaving rodents in the taste field (Chen et al., 2021; Levitan et al., 2019; Fletcher et al., 2017; Roussin et al., 2012), we relied on a single concentration of tastants. Thus we have not tested whether the results generalize across taste intensities. For example, recent studies have shown that a subpopulation of neurons in the rat’s GC (Fonseca et al., 2018) and nucleus accumbens (Villavicencio et al., 2018) track sucrose concentration by increasing their spike coherence with licking. Future work in active licking mice investigating the relative contribution of rate and phase GC coding in representing taste intensity are warranted. In addition, GC neurons are heterogeneous as they have been shown to exhibit layer and cell type specific taste responses (Dikecligil et al., 2020) and are often multimodal (i.e., capable of responding to non-gustatory cues that predict or are contingent upon the availability of food) (Vincis and Fontanini, 2016; Samuelsen et al., 2012; Livneh et al., 2017; Chen et al., 2021). Thus, it might be important to examine whether the GC neurons that encode taste information with a combination of rate and phase coding differ in their identity/function profile compared to the ones in which rate code dominates. The experimental design of the current work did not allow this level of analysis. Future studies will require the use of a more complex experimental design and behavioral task to better understand the observed coding effects in the context of known GC neuron identity and function.

Our data analysis employed a support vector machine learning technique for data classification combined with elastic shape analysis that is designed for data registration (Srivastava and Klassen, 2016). In the context of spike trains, this registration process naturally extracts an alignment function that contains spike timing information (Lu et al., 2014). By using these functions as input to the SVM classification algorithm, one uses timings from the entire post-taste spike trains rather than the less information-rich mean timing. In cases where the timing of spikes in two spike trains are clearly different and stereotyped, averaging over interlick intervals (see Rate-Phase classification described in Fig. 7 and employed in Fig. 8) is an effective way to quantify rate and phase coding. However, if spike trains are not as clearly distinct or stereotyped, then the application of ESA is much more reliable. For example, Fig. 12B shows a raster plot of data from a coding neuron over two interlick intervals and in response to two tastants, salt and sucrose. The neuron fired more near the end of the first interlick interval in response to salt, but fired mostly near the beginning in response to sucrose. This pattern is not apparent during the second interlick interval, when the neuron fired more uniformly in response to salt and at the end of the interlick interval in response to sucrose. There are clear differences in the spiking patterns, but they are far from stereotyped across interlick intervals. However, with the ESA analysis, this neuron performed well when alignment functions were used in the SVM classification (SVM classification rate was 89.3% when using alignment functions). The raster plot of Fig. 12B also shows a clear difference in spike rate during the first interlick interval, which is not so clear in the second interlick interval. The classification score using aligned spike trains picks up the substantial difference in interlick spike rate in response to the two tastants; the neuron has a classification score of 96.3% when aligned spike trains were used in the classification. Overall, the rate/phase quantification obtained using SVM classification with ESA extracts differences in spike trains that are visible by inspection and by using simple techniques like Rate-Phase analysis, but also extracts rate/phase differences that are much harder to see through visual inspection or averaging-based approaches. They are therefore useful tools for separating out rate and temporal coding performed by neurons.

In conclusion, our experiments and analyses quantify the extent by which rate and temporal information can discriminate among tastes in GC neurons, and demonstrate that the timing of licks provides information that can enhance taste discrimination and can therefore be an integral part of the rodent taste experience.

